# Network-level dynamics underlie stress memory and shape the intrinsic limits of mechanically induced defense priming

**DOI:** 10.1101/2024.05.03.592392

**Authors:** Khaoula Hadj-Amor, Ophélie Léger, Sylvain Raffaele, Magdalena Kovacic, Aroune Duclos, Pedro Carvalho-Silva, Frédérick Garcia, Adelin Barbacci

## Abstract

Repeated environmental stimuli can induce transient immune priming in plants, but how such responses accumulate, persist, and decay remains poorly understood. Here, we used acoustic waves to deliver a controlled oscillatory mechanical stimulus and investigated its effects on quantitative disease resistance in *Arabidopsis thaliana*. Repeated stimulations progressively increased resistance to *Sclerotinia sclerotiorum* before reaching a plateau after three stimulations. Transcriptomic analyses showed that priming was not only associated with sustained activation of a restricted set of defense genes, but with progressive reconfiguration of a broad transcriptional response and increased responsiveness to infection. We developed a dynamic network model integrating transcriptional trajectories across repeated stimulations. Modal analysis revealed a distributed dynamical memory composed of rapidly dissipating and slowly relaxing components, providing a basis for the accumulation and transient persistence of priming. The model predicted resistance saturation and rapid relaxation after stimulation ceased, which we experimentally validated. These results support a model in which stress memory is an emergent, multiscale property of transcriptional network dynamics, arising not only from persistent molecular states encoded by a restricted set of genes, but also from their collective dynamics.

## Introduction

Plants have evolved a complex immune system achieved through interconnected, layered molecular interactions (Roux *et al*., 2014; Delplace *et al*., 2020; Remick *et al*., 2023). The perception of conserved pathogen-associated molecular patterns (PAMPs) by surface Pattern Recognition Receptors (PRRs) activates PAMP-triggered immunity (PTI). Pathogens may deploy effectors that suppress PTI or activate a second layer known as Effector-Triggered Immunity (ETI) upon recognition by intracellular receptors. Both PTI and ETI operate through receptor-based molecular lock-and-key mechanisms that can confer strong or complete resistance when recognition occurs. However, minor variations in effector structure may disrupt this specificity (Raffaele *et al*., 2010; Remick *et al*., 2023; Derbyshire & Raffaele, 2023), and necrotrophic pathogens that thrive on dead plant cells can exploit programmed cell death to promote infection (Govrin & Levine, 2000; Barbacci *et al*., 2020; Derbyshire & Raffaele, 2023).

In the absence of such complete immunity, a form of disease resistance known as quantitative disease resistance (QDR) is observed, which mitigates rather than halts disease progression (Poland *et al*., 2009; Wang *et al*., 2019). QDR frequently occurs in response to necrotrophic fungi such as *Botrytis cinerea* or *Sclerotinia sclerotiorum*. While QDR can incorporate receptor-based key–lock interactions similar to those underlying PTI and ETI, it is fundamentally a highly multigenic process involving large-scale regulatory reprogramming across thousands of genes, extending beyond the sparse recognition events characteristic of PTI/ETI. Transcriptomic analyses of six species within the Pentapetalae clade revealed that approximately 30% of plant transcriptomes respond locally to infection by *S. sclerotiorum* (Sucher *et al*., 2020). Consequently, genes associated with QDR tend to be pleiotropic and not specific to the immune response, a pattern consistent with neutral network rewiring of transcriptomes observed across *Arabidopsis* accessions (Delplace *et al*., 2025), in which abundant regulatory variation can be integrated without loss of resistance. The highly multigenic nature of QDR makes it prone to influence by abiotic environmental stresses, such as temperature variations (Aoun *et al*., 2017; Singh *et al*., 2023; Didelon *et al*., 2026), as well as internal cues such as mechanical fluctuations in cell walls (Engelsdorf *et al*., 2018), which may be indirectly induced by necrotrophic fungi (Léger *et al*., 2022).

Prior exposure to a mild stress can condition plants to respond more effectively to a subsequent, more severe stress of the same type, resulting in an enhanced response upon repeated exposure (Wang *et al*., 2014; Baier *et al*., 2019). For example, plant defense mechanisms can be activated by exposing plants to avirulent pathogens, thereby conferring protection against secondary infections (Verhagen *et al*., 2010). This phenomenon, in which a stress response is exacerbated by prior exposure, is often referred to as “priming” (Conrath, 2011; Liu *et al*., 2022). Stress exposure may also impart tolerance to unrelated stresses, as observed in wheat plants exposed to mild drought conditions that subsequently demonstrate increased resistance to low temperatures (Ru *et al*., 2024). This is typically designated as “cross-priming” and may rely on the potentiation of responses distinct from those triggered by the initial stress. Priming is thought to involve forms of stress memory that allow information acquired during a previous exposure to influence subsequent responses, although the mechanisms underlying such memory remain incompletely understood (Bruce *et al*., 2007; Crisp *et al*., 2016). Through alterations in morphology, physiology, transcriptional and epigenetic regulation, and other cellular processes, priming can enhance plant adaptation to fluctuating environments. The robustness of priming phenomena may therefore depend not only on the mechanisms activated by an individual stress exposure, but also on how information from successive exposures is integrated and retained over time. Understanding how such memory is established, maintained, and dissipated is thus important both for understanding plant adaptation and for exploiting priming in sustainable crop protection (Liu *et al*., 2022).

The robustness of priming phenomena in fluctuating environments depends on the temporal dynamics and breadth of the underlying stress memory mechanisms. Memory can be viewed as a dynamic process through which information acquired during a previous exposure influences subsequent states and responses. In plants, memory can take a myriad of forms characterized by distinct durations (Thellier *et al*., 2018). On a temporal scale of hours, rudimentary memory mechanisms, exemplified by the homogalacturonan cycle linked to pressure fluctuations, appear to operate autonomously from genetic regulation (Proseus & Boyer, 2006; Barbacci *et al*., 2013). Over time spans ranging from hours to generations, transcriptional memory involving transcription factors and epigenetic modifications has emerged as a prominent stress memory mechanism (Avramova, 2019). For example, trimethylation of histone H3 at lysines 4 and 9 (H3K4 and H3K9) is a major epigenetic modification associated with the acclimation of poplar trees to recurrent mechanical stimuli. A single bending event is sufficient to induce significant alterations in the abundance of these epigenetic marks (Ghosh *et al*., 2023). Transcription factors, which regulate gene expression and mediate signaling pathways, also play crucial roles in transcriptional memory, operating in synergy with epigenetic modifications. A paradigmatic example is FORGETTER3/HEAT SHOCK TRANSCRIPTION FACTOR A3 (FGT3/HSFA3), which is required for the establishment of physiological heat stress memory and for maintaining increased expression of memory-related genes in the days following heat exposure (Friedrich *et al*., 2021). While these mechanisms provide important molecular substrates for stress memory, they do not fully explain how information from repeated exposures is coordinated across the transcriptome and integrated over time. In particular, locus-specific transcriptional or epigenetic changes may represent components of broader regulatory dynamics rather than independent memory switches. This raises the possibility that stress memory can emerge from the collective dynamics of interacting regulatory networks, in which successive stimuli progressively reshape the transcriptional state of the plant. Such a framework could explain how memory accumulates across repeated exposures while remaining reversible, and could also impose intrinsic constraints on the rate of memory formation and its persistence after stimulation has ceased. Within this view, priming may reflect a form of network-encoded stress memory arising from the dynamic reconfiguration of gene regulatory interactions rather than from the persistent activation of a restricted set of genes.

To investigate how repeated environmental stimuli can be integrated into stress memory at the network level, we used acoustic stimulation as a controlled mechanical stimulus. Sound, characterized by minute temporal fluctuations in pressure, represents a form of mechanical stimulation. Although sound perception mechanisms remain unclear, numerous effects on plant biology have been documented (Appel & Cocroft, 2014, 2023; Hassanien *et al*., 2014; Demey *et al*., 2023). Concerning defense, brief exposure of *Arabidopsis thaliana* to fixed-intensity sound vibrations modulates genes involved in defense responses and signal transduction pathways (Ghosh *et al*., 2016). The affected genes depend on sound frequency, yet several disease-resistance–associated genes, including AtHSPRO2 (At2g40000), AtJAZ7 (At2g34600), and At5g66070, respond to multiple frequencies. Notably, phenotypic gains in disease resistance are observed only after repeated acoustic stimulation (AS), with *A. thaliana* becoming more resistant to *B. cinerea* after 10 days of daily sound exposure (Ghosh *et al*., 2017). Transcriptomic and proteomic analyses revealed a profound effect of repeated AS on the reprogramming of primary metabolism. However, the system-level processes through which repeated AS reshapes plant regulatory states and promotes a transition from susceptibility to resistance remain unknown.

To gain insights into the architecture and dynamics of stress memory underlying priming, we investigated how repeated acoustic stimulation reshapes transcriptional states over time and how these changes relate to the dynamics and persistence of defense priming against *Sclerotinia sclerotiorum*. By integrating genetic lines impaired in mechanical perception (mca1 and msl4/msl5/msl6/msl9/msl10), genome-scale transcriptomics, and dynamic network modeling, we asked whether the cumulative properties of acoustic-induced resistance could emerge from the collective dynamics of the transcriptional network. We first characterized how resistance and transcriptional states changed with increasing numbers of acoustic stimulations, and then inferred a dynamic network model to determine how these trajectories could be decomposed into dynamical components with distinct persistence. We further tested whether the resulting network dynamics could account for experimentally observed constraints on the accumulation and persistence of priming, including the effects of stimulation frequency and interruption. Together, these approaches allowed us to examine stress memory as a distributed property of network-level regulation, while testing whether the dynamics of the inferred network impose quantitative limits on the formation and persistence of the primed state.

## Material and Methods

### Plant/Fungal material cultivation and inoculation

Arabidopsis plants were cultivated under constant conditions, including a 9-hour light period at 120 µmol/m²/s, a temperature of 22°C, and a relative humidity of 70% for 4 weeks before inoculation and RNA sampling. The *Sclerotinia sclerotiorum* isolate (1980) was cultured on potato dextrose agar (PDA) plates at 25°C for 3 days before inoculation. Inoculation was carried out by placing a 5 mm diameter PDA plug on the adaxial surface of *A. thaliana* leaves, following the methodology described in Barbacci *et al*., (2020). The mock treatment involved using an *S. sclerotiorum*-free plug. For phenotypic assessments, inoculations were conducted on detached leaves within a humidity-controlled chamber, facilitating disease progression imaging. RNA-seq analyses were performed on whole plants subjected to inoculations in the same chambers.

### Phenotyping

Phenotyping was conducted on detached leaves within a controlled environment characterized by elevated humidity and a constant temperature. The enclosed space, which was equipped with a camera, captured images at 10-minute intervals. Analysis of these images allowed the measurement of lesion sizes on each leaf during infection. The susceptibility of individual leaves was subsequently calculated as the growth rate during the exponential phase of disease development, employing the segmented package in R (R Core Team, 2021). Each phenotyping analysis presented in this paper was performed independently and repeated at least three times. For more information, please consult Barbacci *et al*., (2020). The Python code for image analysis is available at https://github.com/darcyabjones/INFEST.

### Sound priming and RNA sampling

After three weeks, the plants were transferred to two growth chambers under identical cultivation conditions. One growth chamber was equipped with a speaker (4” Visaton full range) connected to a power amplifier. The plants were subjected to acoustic stimulations during the initial three hours of the day (or the first six hours of two daily stimulations). The acoustic waves utilized were 1 kHz sinusoidal waves at a 100 dB intensity. This frequency and intensity were selected based on prior evidence that 1000 Hz sinusoidal waves at 100 dB induce significant transcriptional changes in defense-related genes in Arabidopsis (Ghosh *et al*., 2016, 2017). The 100 dB intensity was selected as a calibrated mechanical stress to systematically probe transcriptional memory constraints while ensuring the stimulus substantially exceeded growth chamber background noise (∼70 dB). This high-intensity regime was chosen as a calibrated mechanical stress to systematically probe the upper boundaries and dynamic constraints of transcriptional memory, rather than to mimic specific ecological conditions. Acoustic intensity was measured at the top of the plants via a calibrated sound level meter. Plants and leaves did not exhibit obvious variations in development after AS stimulation (plant or leaf size, color, etc.). Infected plants were inoculated at the end of the final AS session. Leaf samples were collected after the last acoustic stimulation for non-inoculated plants and 24 hours post-inoculation (hpi) for infected plants. Leaves were cut with a razor blade and immediately placed on a glass cooled with liquid nitrogen as described in Sucher *et al*., (2020). For every plant, a 2 mm thick ring around the infected area or the *S. sclerotiorum*-free plug (mock) was taken for RNA extraction.

### RNA extraction and differential gene analysis

RNA extraction was performed via the Nucleospin RNA kit from Macherey-Nagel. Subsequently, cDNA synthesis was carried out employing Superscript III reverse transcriptase (Invitrogen Carlsbad, CA) and 1 μg of total RNA. For RNA-seq, reads were concurrently mapped against the genomes of *A. thaliana* Col-0 (TAIR10 release) and *S. sclerotiorum* 1980. This was accomplished using the nf-core/rnaseq pipeline version 3.0 (Patel *et al*., 2020), with specific parameters, namely “–– skip_alignment –-pseudo_aligner salmon.” This set of parameters included adapter and quality trimming using Trim Galore software version 0.6.6, followed by transcript assignment and quantification using the salmon tool (version 1.4.0).

### Differential gene analysis

Three differential gene expression analyses were conducted. To elucidate the impact of repeated acoustic stimulation (AS) on gene expression in non-inoculated plants, we utilized the gene expression levels of uninfected plants that were not exposed to acoustic stimulation as a reference. Differentially expressed genes for non-inoculated plants exposed to 1, 3, and 8 ASs were then identified through three paired statistical tests based on the negative binomial distribution. Gene expression levels associated with Bonferroni-adjusted p-values less than 0.05 and with ||log2 fold change|| > 2 were retained for further analysis. The same analytical process was employed to identify differentially expressed genes between consecutive ASs in non-inoculated plants and to understand the ASs effect on plants exposed to acoustic stimulation before inoculation. In the former case, the reference was the gene expression level of plants exposed to the previous ASs number. In the latter case, the reference was the gene expression level of plants not exposed to acoustic stimulation, considering infection as the reference. The analysis of differentially expressed genes was conducted via the R package DESeq2 (Love *et al*., 2014).

### Data analysis

Enrichment analysis were performed via TopGO (Alexa *et al*., 2006). P-values are obtained via the hypergeometric test, whereas false discovery rates (FDRs) are adjusted through the Benjamini-Hochberg method to account for multiple tests. Fold enrichment is calculated as the ratio of the percentage of genes in the given list associated with a pathway to the corresponding percentage in the background gene set. While FDR is used to assess statistical significance, fold enrichment is used to quantify the effect size. Enrichments were calculated relative to a reference background. For Figures 1C and 4E, the background consisted of the set of differentially expressed genes for the previous number of AS. As shown in Fig. 1C, the plants were not inoculated, whereas as shown in Fig. 4E, the plants were inoculated. For the enrichments of the clusters shown in Sfig. 1, enrichments were calculated relative to the set of all genes present in the 25 clusters. Transcription factors and transcription factor regulations were collected from the public database PlantTFDB (Jin *et al*., 2017) and are available in **SFile 4**. Statistics were computed using R software (R Core Team, 2021). The test used is indicated in the figure legend and the main text. Plots were made with ggplot2 (Wickham, 2009) and circlize (Gu *et al*., 2014) R libraries and combined with Inkscape.

**Figure 1.**
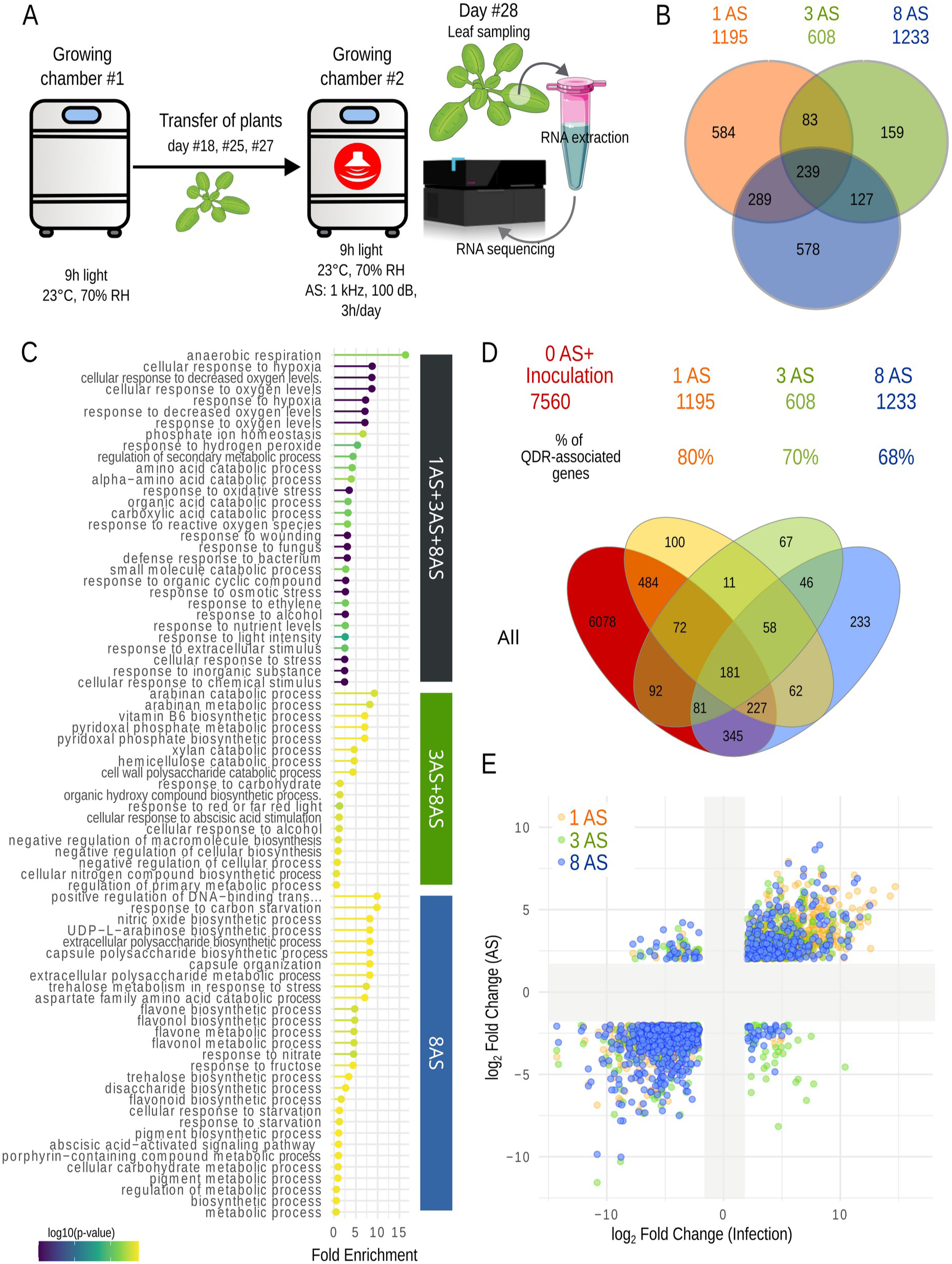
Impact of repeated acoustic stimulations (AS, 3 h/day, 1000 Hz, 100 dB) on gene expression in non-inoculated plants. **A.** Overview of the experimental protocol. **B.** Venn diagrams illustrating DEGs after 1, 3, and 8 AS in non-inoculated plants. Numbers indicate the total DEGs in each set. **C.** Enrichment analysis of biological process ontologies over sequential AS. Colors indicate –log10(p-value) for each term. **D.** Venn diagrams showing DEGs in non-inoculated plants after 1, 3, and 8 AS compared with DEGs in plants inoculated with *Sclerotinia sclerotiorum* without AS. Numbers indicate the total DEGs in each set. **E.** Log2 fold changes of DEGs in response to infection and 1, 3, or 8 AS. Gray shading denotes the selection threshold of ||log2 fold change|| > 2.

### Clustering and dynamic gene network inference

The dataset details the expression profiles of 2059 genes that were differentially expressed across four distinct time points. The dimensions of the dataset initially posed a challenge for network inference. To address this, a crucial first step involved the substantial reduction of data dimensions using a clustering method. Traditional clustering approaches designed for static or dynamic data, such as k-means, were deemed unsuitable for our sparsely distributed temporal data. Consequently, to categorize very short time series characterized by irregularly spaced time points, we performed differential gene analysis between consecutive time measurements to elucidate the temporal variations in each gene and grouped genes with similar trajectories. Where expression significantly differed between two successive time points, the variations were designated as either positive (+) or negative (-). If the differences were not statistically significant, the temporal variation was denoted as zero (=). Subsequently, genes were organized into 27 clusters based on similar expression signatures. Notably, the cluster representing genes with a constant expression (===) was excluded from further analysis. The dynamic control network was derived through computation based on an extended explicit difference equation model, formulated as follows: X_N+1_=A.X_N_+ AS_Systematic effect_. Here, X_N_, X_N+1_, and the AS_Systematic effect_ are vectors of length equal to the cluster number K=25, and A is a K×K square matrix. This equation represents the variation in the expression of a cluster-representative gene over time as a linear combination of the expression of cluster-representative genes of all clusters and a constant term AS_Systematic effect_. The fixed (systematic) effect of sound on cluster-representative gene expression is described by AS_Systematic effect_. The model was constrained by the condition A.X_0_=0, reflecting the fact that AS starts on the first day from an initial condition that we assume to be in equilibrium relative to the dynamics described by the matrix A. We also set AS_Systematic effect_ = 0 for clusters whose first transition between N=0 and N=1 was of the “=” type. To estimate A and AS_Systematic effect_ from the previous equation, we first imputed missing data along the trajectories at a time step of one day, using simple linear interpolation. The estimation of A and B from the experimental data was performed cluster by cluster, that is, row by row for A and B, using the Elastic Net regression method, implemented in the R package glmnet (Friedman *et al*., 2010). We evaluated several approaches to define representative cluster expression values X_N_ (including mean values, quantiles, and randomly sampled gene expressions) for predicting X_N+1_ using linear models. The best predictive performance, assessed by mean squared error, was obtained using a gene-level approach. Specifically, for each gene g within a given cluster, we modeled the expression change X_g,N+1_−X_g,N_ as a linear function of its own expression X_g,N_ and the expression levels of the K−1 other clusters, represented by the quantile corresponding to gene g within each cluster. Model parameters were estimated using elastic net regularization, with weights of 0.99 and 0.01 assigned to the Lasso (Tibshirani, 1996) and Ridge penalties, respectively, and 30-fold cross-validation.

### Modal analysis of network dynamics

To investigate transcriptional memory induced by repeated acoustic stimuli (AS) in *Arabidopsis thaliana*, we carried out an initial modal analysis of the autonomous dynamic system defined by the equation X_N+1_ = A X_n_, where A is the inferred regulation matrix.

The matrix A has 17 distinct eigenvalues, including 7 real roots and 10 conjugate double roots. The eigenvalue 1 has order 9, but the dimension of the associated eigenspace (the kernel of A) is only 8. The matrix A is therefore not diagonalizable in a basis of eigenvectors. It is, however, possible to supplement the 8 eigenvectors associated with eigenvalue 1 with a 9th generalized eigenvector derived from the kernel of A², according to the principle of generalized eigenspace decomposition (Weintraub 2008).

We thus constructed a new basis consisting of the 24 original eigenvectors and this generalised eigenvector, in which the matrix A is diagonal except for the last column. By grouping the complex eigenvalues in pairs as usual, these 25 basis vectors define 12 generalized modes of the system’s dynamics, called metamodes. These metamodes can be analysed as in the classical diagonalizable case, keeping in mind, however, that the presence of the 25th generalised eigenvector results in coupling between this mode and the others.

The contribution of each metamode to the transcriptional dynamics across repeated stimulations can be quantified by the norm of the projection of the transcriptional states X_N_ onto the corresponding eigenspace. This allows the characterisation of how individual metamodes are activated and maintained over time.

The magnitude of the eigenvalue |λ| of a metamode provides information about its general intrinsic dynamics. Modes whose eigenvalue magnitude is strictly below 1 correspond to stable components that decay following a perturbation. The characteristic decay time is then defined by τ = −1/log(|λ|), and a time t = 3τ is taken to represent a return to 95% of the mode’s equilibrium value.

## Results

### Acoustic stimulation pre-activates multiple defense pathways prior to pathogen exposure

To document the molecular changes induced by the repetition of acoustic stimulation (AS), we analyzed the global gene expression of 4-week-old *A. thaliana* Col-0 plants exposed to AS corresponding to 3 hours of 1000 Hz sinusoidal waves at 100 dB (∼2 Pa) once a day for 1, 3 or 8 days (**Fig. 1A**). A mature leaf from each plant was then collected for RNA extraction. Compared with plants not exposed to AS, approximately 1000 *Arabidopsis thaliana* genes were differentially expressed (DEGs, ||log_2_FC|| > 2 and p-value < 0.05) in response to each AS treatment (**Fig. 1B, SFile 1**). Overall, 8% of the *A. thaliana* genes (2059) were DEGs under at least one AS treatment, among which 239 were DEGs independent of the number of AS (in all treatments), and 1820 were modulated by a specific number of AS. This temporal transcriptomic reprogramming across increasing numbers of acoustic stimulations (AS) indicates that, beyond a core set of genes consistently responsive to AS, a large proportion of differentially expressed genes exhibits expression patterns that depend on stimulation number, suggesting transcriptional features consistent with memory.

To highlight how plants implement their responses over repeated stimulations, we analyzed enriched gene ontologies (GOs) among up-regulated DEGs from 1 AS (DEGs after 1, 3 and 8 AS), from 3 AS (DEGs after 3 and 8 AS but not 1 AS), and DEGs specific to eight AS (**Fig. 1C, SFile 2**). From the very first stimulation, the analysis of enriched GO terms suggested that plants may rapidly coordinate metabolic and immune responses by modulating processes related to hypoxia and oxidative stress, secondary metabolism, and immune defenses against fungi and bacteria (**Fig. 1C**). Starting from the third stimulation, the ontological analysis indicated a possible shift toward cell wall remodeling, metabolic fine-tuning, and hormone-mediated defense, notably through the catabolism of hemicellulosic polysaccharides and the modulation of vitamin B6 pathways involved in stress adaptation, including pyridoxal phosphate biosynthesis and abscisic acid signaling (**Fig. 1C**). At the eighth stimulation, the enrichment of additional GO terms proposed pronounced metabolic reprogramming, enhanced stress adaptation, and reinforced immune responses, exemplified by the activation of abscisic acid signaling, the biosynthesis of extracellular polysaccharides, flavone/flavonol metabolic processes, and broad regulation of metabolic pathways (**Fig. 1C**). Among 31 epigenetic regulators previously associated with thermomemory and defense (Ding & Wang, 2015; Liu *et al*., 2022), only four exhibited AS-dependent expression patterns (**SFile 3**). While this suggests that large-scale transcriptional induction of canonical epigenetic machinery is not the primary driver here, it does not rule out the involvement of post-translational activations or localized chromatin remodeling. Therefore, the precise contribution of baseline epigenetic factors during acoustic priming remains an open question that warrants dedicated epigenomic investigations. More broadly, rather than pointing to a persistent activation of specific individual pathways, these results demonstrate that acoustic memory is highly dynamic. They support the view that the transcriptional memory of AS did not result from the sustained activation of mechanisms triggered after the first AS, but rather from the activation of parallel mechanisms during repeated AS treatments.

To investigate how AS-induced transcriptional reprogramming affects plant immune responses, we compared AS-triggered DEGs with those from plants inoculated with the necrotrophic fungus *Sclerotinia sclerotiorum*. Enrichment analysis suggested a progressive establishment of the immune response, which was confirmed by comparing transcriptomes of plants after repeated AS with plants not exposed to AS 24 hours post-inoculation with *S. sclerotiorum* (**Fig. 1D**). Among the 7560 DEGs identified upon fungal inoculation, 1482 (20%) were also differentially expressed in response to AS alone. Notably, 70% of DEGs after three AS treatments (608 genes) and over 68% after one or eight AS treatments (1195 and 1233 genes, respectively) overlapped with pathogen-responsive DEGs, whereas only 181 genes were modulated upon pathogen inoculation and regardless of the number of AS (**Fig. 1D**). Gene expression changes induced by repeated AS occurred in the same direction as those observed in inoculated plants for over 84% of genes, although the average log2 fold change was roughly half that observed in pathogen-inoculated plants (**Fig. 1E**). Thus, a large proportion of AS-responsive genes were also associated with the plant response to *S. sclerotiorum*. Together, these results indicated that repeated AS was associated with a progressive, stimulation-dependent engagement of regulatory pathways linked to pathogen responses, consistent with the emergence of memory-like behavior at the level of large-scale transcriptional reprogramming.

### Sequential recruitment of transcription factor targets diversifies AS-mediated transcriptome reprogramming at each stimulation

Consistent with the likely limited involvement of canonical epigenetic regulators, we hypothesized that the diversification of biological processes observed across successive AS treatments could arise from the progressive activation of transcription factors (TFs) with distinct target gene repertoires. To explore this possibility, we analyzed TFs differentially expressed upon AS and their predicted target genes, based on cis-regulatory motifs curated in PlantTFDB (Jin *et al*., 2017) (**Fig. 2**, **SFile 4**). TF expression was strongly dependent on the number of AS exposures, with only 9 TFs (12%) being modulated independently of AS repetition. After a single AS, 62 TFs (TFs_1AS_) were differentially expressed, together with 817 of their predicted target genes, representing 68% of all DEGs identified at this stage. Notably, among the predicted targets of TFs_1AS_, 37 were TFs that became differentially expressed only after three AS treatments (TFs_3AS_). At three AS, 22 TFs specifically differentially expressed at this stage were associated with 258 DEGs, corresponding to 42% of all DEGs. Among their predicted targets, 15 TFs were differentially expressed only after eight AS treatments (TFs_8AS_). At eight AS, 34 TFs were associated with 750 DEGs, accounting for 61% of all DEGs at this stage. Importantly, both the identity and number of TFs engaged varied across stages, despite partial overlap in predicted regulatory relationships. This pattern suggests that TFs activated at earlier stages contribute to progressive transcriptional remodeling, thereby shaping regulatory interactions observed after subsequent AS exposures. Overall, 67% of repeated AS-responsive DEGs (2059 genes) could be associated with predicted targets of TFs sequentially recruited across AS repetitions. Although not demonstrating causality, this analysis provides a framework for the temporal dynamics of AS-induced transcriptional changes (**Fig. 2**). In this context, and by analogy with mitotic bookmarking described in animals (Bellec *et al*., 2018), our results were consistent with the notion that the progressive engagement of TF-centered regulatory networks may contribute to the persistence of prior transcriptional states.

**Figure 2.**
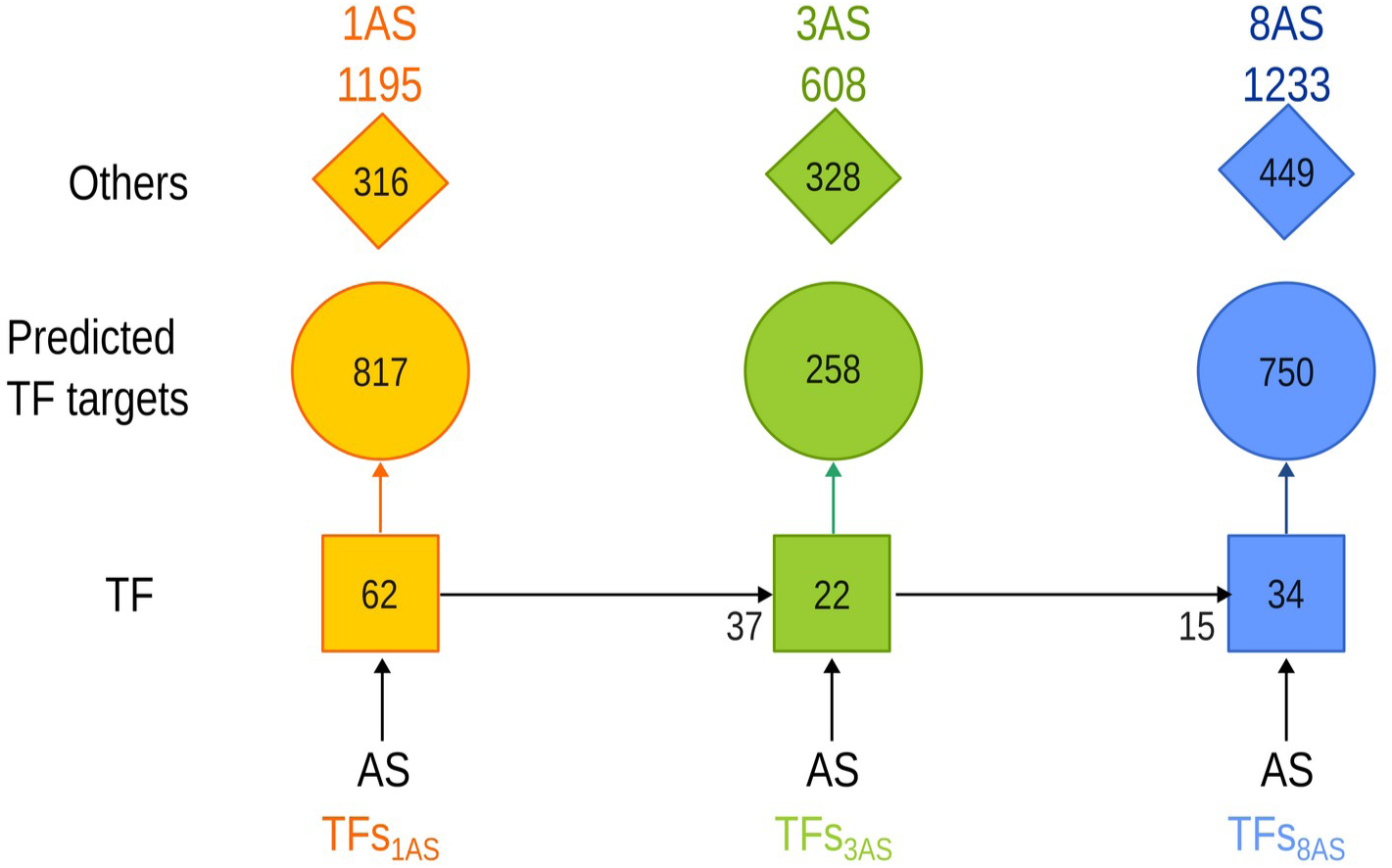
Transcription factor regulatory cascade across repeated acoustic stimulations (AS). Squares represent transcription factors (TFs), with numbers indicating the number of predicted target DEGs at each AS stage. Numbers adjacent to squares denote TFs predicted to be regulated by TFs activated at the previous stage. Circles represent non-TF DEGs, and diamonds indicate AS-responsive DEGs not assigned to any TF in the network.

### AS-induced disease resistance requires repeated stimulations

To investigate the impact of AS on plant defense, we applied AS corresponding to 1000 Hz sinusoidal waves at 100 dB for 3 hours every day for 0, 1, 2, 3, or 10 days before inoculation with *S. sclerotiorum* (**Fig. 3A, SFile 5**). Disease resistance was quantified by evaluating the growth rate of necrosis in leaves according to Barbacci *et al*., (2020). A single AS led to a 16% increase in disease susceptibility (Wilcoxon rank-sum test p-value 4.3E-4), whereas a minimum of 2 AS led to an increase in disease resistance. Two AS elicited a slight but significant 11% increase in disease resistance (Wilcoxon rank-sum test p-value 0.022). Beyond 3 AS, a substantial ∼25% gain in disease resistance was observed and remained similar in plants subjected to 10 AS (Wilcoxon rank-sum, p-value 3.3E-5 and 1.2E-5 at 3 and 10 AS respectively, **Fig. 3A,D**). This AS-induced resistance was largely independent of trichome morphology or density, as a similar resistance gain after 10 AS was observed in glabrous mutant plants lacking trichomes (*gl-1*,Benikhlef *et al*., 2013) (Wilcoxon rank-sum test p-value: Col-0 0.003, *gl-1*: 0.0007 **Fig. 3B, SFile 6**). To further evaluate the robustness of AS-induced responses, we analyzed *mca1* (Yamanaka *et al*., 2010) and the *Δmsl5* (*msl4/msl5/msl6/msl9/msl10*) quintuple mutant line (Tran *et al*., 2021), two mutants impaired in mechanoperception and displaying increased susceptibility to *S. sclerotiorum* (**Fig. 3C, SFile 7**). Because mechanoperception contributes to quantitative disease resistance (Léger *et al*., 2022), these mutants allowed us to test whether AS responses result primarily from signal perception or network-level integration contributing to stress memory. After 3 daily AS, disease resistance in these mutants was similar to wild-type levels, suggesting that repeated AS progressively restores resistance through compensatory regulatory mechanisms (**Fig. 3C**). Together, these results indicate that AS-induced resistance depends on temporal integration of repeated stimuli and can partially compensate for impaired mechanoperception.

**Figure 3.**
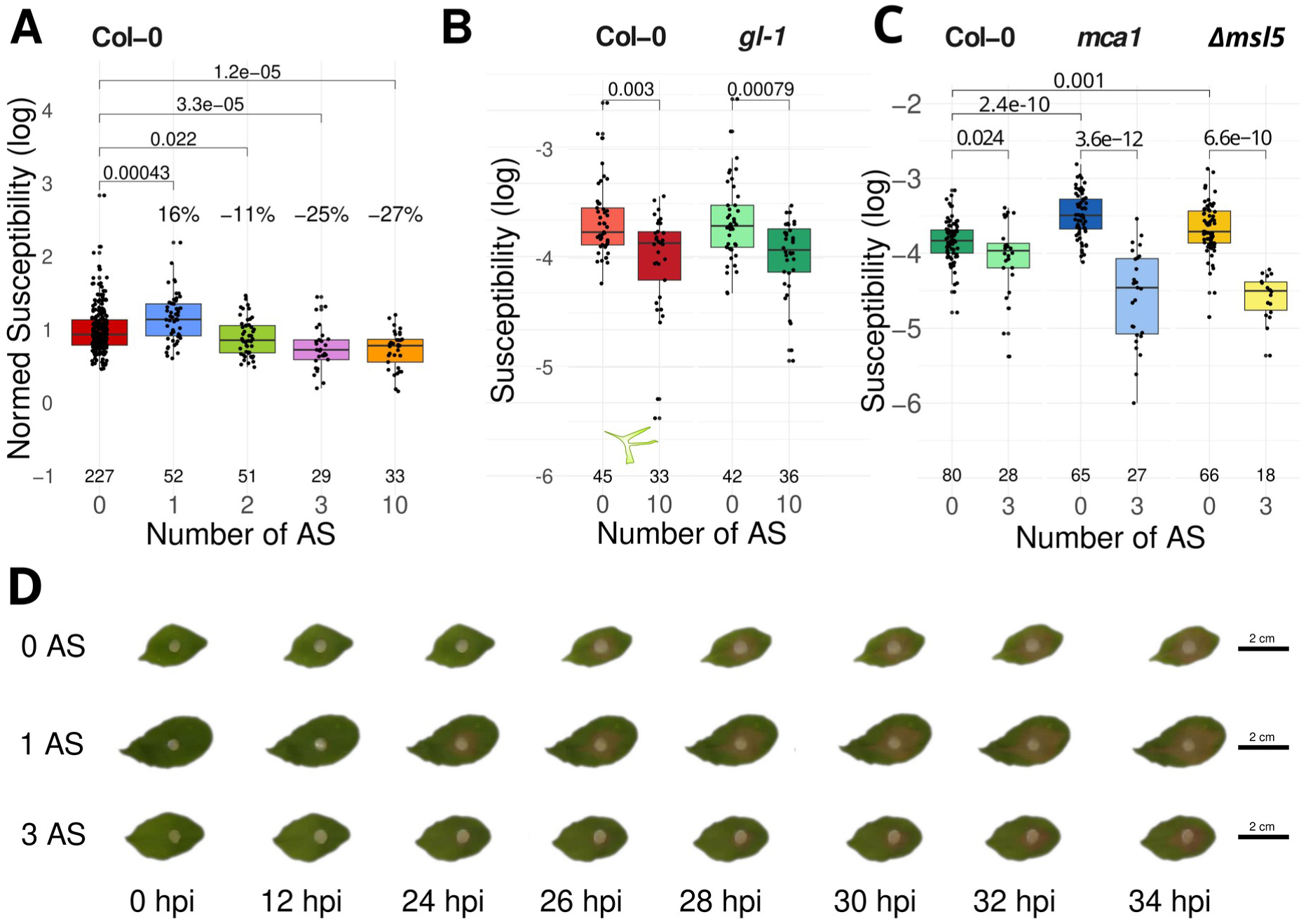
Effect of repeated acoustic stimulations (AS) on plant susceptibility to *Sclerotinia sclerotiorum*. **A.** Changes in susceptibility with increasing numbers of AS. **B.** Susceptibility of wild-type and trichome-lacking genotypes across different numbers of AS. **C.** Susceptibility of the *Δmsl5* quintuple mutant and *mca1* mutant lines following exposure to AS. Boxplots represent the interquartile range (25th–75th percentile) with the median indicated by the bold line. Individual data points are overlaid using jitter for clarity. Unpaired Wilcoxon rank sum test p-values are shown, and sample sizes are indicated below each boxplot. **D.** Representative progression of lesion development caused by *Sclerotinia sclerotiorum* on Col-0 *A. thaliana* leaves over 34 hours post-inoculation (hpi), shown for plants not exposed to AS (0 AS), and for plants exposed to 1 or 3 AS treatments.

### Repeated AS broadened and intensified the activation of QDR-associated genes to prime plant defense

To investigate transcriptional changes associated with disease resistance priming by repeated AS, *A. thaliana* Col-0 plants were exposed to identical AS (1000 Hz, 100 dB, 3 h) once per day for 0, 1, 3, or 8 consecutive days prior to inoculation with *S. sclerotiorum*. RNA sequencing was performed 24 h post-inoculation (**Fig. 4A**). As shown previously, differential expression analysis comparing inoculated plants with non-inoculated naïve plants identified 7560 genes responsive to infection in the absence of prior acoustic stimulation (**Fig. 1D**, **SFile 8**). Of these, 93% were also differentially expressed in plants that had received one or more prior acoustic stimulations before inoculation ( **Fig. 4B**), defining a large core inoculation-responsive transcriptional program consistent with the dominant influence of the most recent stress in sequential stress contexts.

**Figure 4.**
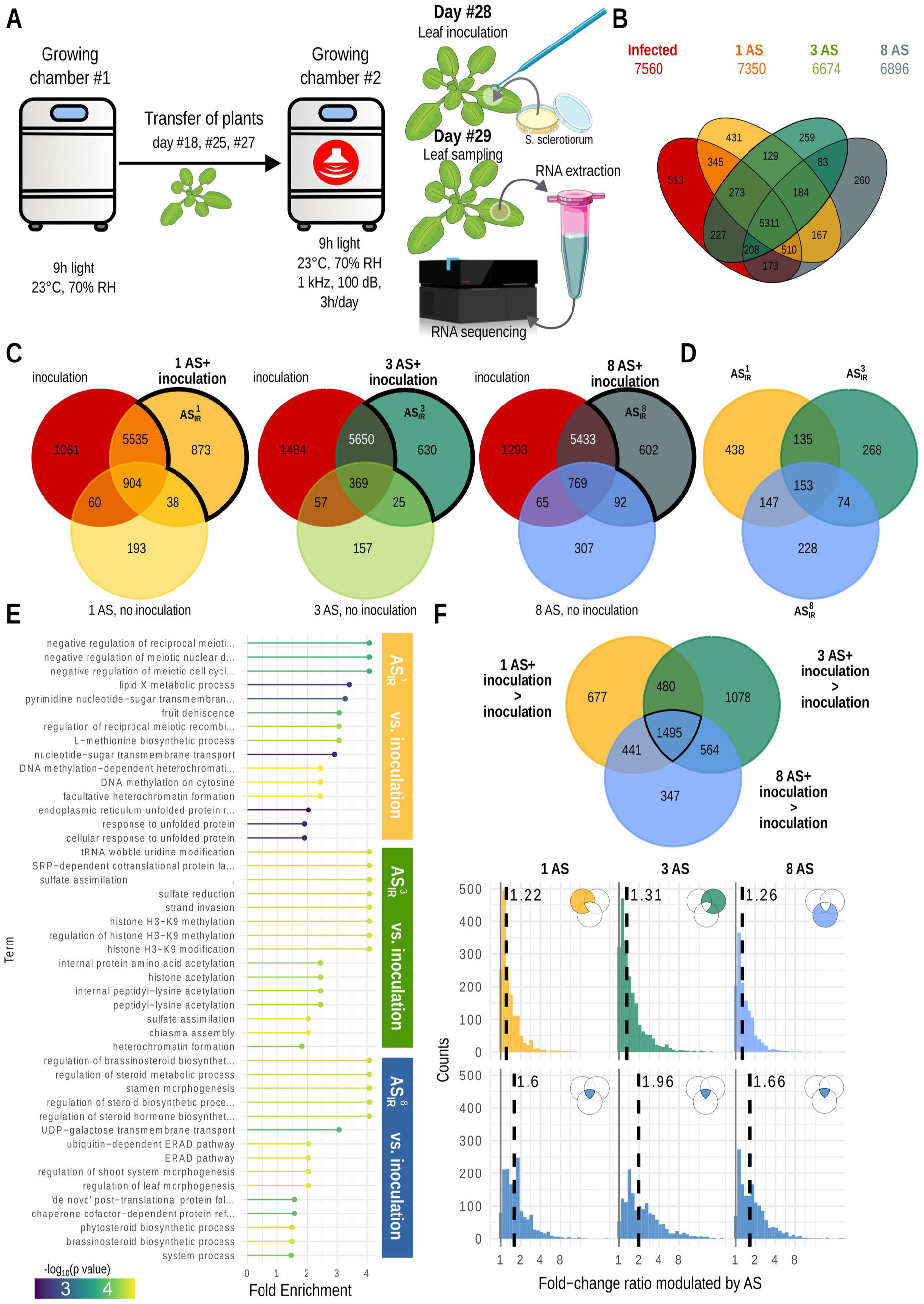
Impact of acoustic stimulation (AS) on the plant transcriptome during subsequent infection. **A.** Schematic overview of the experimental protocol. **B.** Venn diagrams showing the overlap of differentially expressed genes (DEGs) in infected plants previously exposed to 1, 3, or 8 AS treatments, compared with infected naïve plants (control). **C.** Venn diagrams illustrating genes modulated during infection in response to prior AS. Gene expression in infected plants with a history of AS is compared against their respective non-inoculated but AS-exposed controls. ASIR_N denotes the set of AS-dependent, infection-responsive genes after the Nth AS. **D.** Venn diagram highlighting the subset of AS-dependent infection-responsive genes that are already modulated by AS in the context of infection. **E.** Gene Ontology (GO) enrichment analysis for biological processes among AS-dependent, infection-responsive genes. **F.** Venn diagrams and distribution of induction ratios for genes showing enhanced expression upon infection in plants with prior AS exposure compared to infected naïve plants. The top row shows the distribution for genes whose overexpression depends on the number of prior AS, while the bottom row shows genes overexpressed irrespective of AS number. Dashed lines indicate the median of each distribution.

To identify transcriptional responses specifically requiring prior AS, we compared (i) plants exposed to AS but not inoculated, (ii) plants inoculated after prior AS, and (iii) plants inoculated without prior AS (**Fig. 4C**). Genes differentially expressed in AS-treated inoculated plants relative to both naïve inoculated plants and AS-treated non-inoculated plants were defined as AS-dependent inoculation-responsive genes (AS_IR_). We identified 873 AS_IR_ genes after a single prior AS, 630 after three consecutive daily AS, and 602 after eight consecutive daily AS, yielding a total of 1443 AS _IR_ genes. This represented a 19% increase relative to the infection response observed in naïve plants. Only 153 AS_IR_ genes (11%) were shared across all stimulation regimes, indicating that diversification of infection-responsive transcription strongly depends on the number of identical prior AS (**Fig. 4D**).

Gene Ontology enrichment analysis of AS-dependent infection-responsive genes (AS_IR_) compared to gene expressed in naïve infected plants revealed distinct functional signatures depending on the number of prior stimulations (**Fig. 4E**; **SFile 9**). After 1 AS, AS_IR_ genes were enriched in categories related to chromatin organization, protein homeostasis, and metabolic adjustment, suggesting early reconfiguration of regulatory and stress-associated pathways. After three AS, enrichment shifted toward processes associated with translational regulation, sulfur metabolism, and chromatin dynamics, consistent with consolidation of transcriptional and metabolic reprogramming. After eight daily stimulations, AS_IR_ genes were predominantly enriched in steroid biosynthesis, protein quality control, and developmental regulation, indicating integration of hormonal and growth-related pathways under prolonged repetitive stimulation.

We next examined whether prior acoustic stimulation quantitatively enhanced the induction of core inoculation-responsive genes, defined as genes modulated by inoculation both in naïve (no-AS) and AS-treated plants. Among core inoculation-responsive genes, 1495 exhibited AS-independent induction, whereas 3587 showed AS-dependent induction (**Fig. 4F**). For each core gene, we calculated an induction ratio corresponding to expression in stimulated plants divided by expression in naïve after inoculation. Gene expression peaked after three consecutive daily stimulations in both AS-dependent and independent genes, with median induction ratios of 1.96 for the AS-independent group and 1.31 for the AS-dependent group. Together, these results indicated that repeated exposure to an identical acoustic stimulus enhanced inoculation-triggered transcription through a stable core program, complemented by an additional set of genes whose recruitment depended on the number of prior daily stimulations, thereby progressively diversifying the immune response in a stimulation-number-dependent manner.

### Repeated acoustic stimuli establish distributed stress memory through network-level regulation

To gain a better understanding of the regulatory basis of AS-induced stress memory, we investigated how transcriptional responses were organized across successive AS treatments. Given the large number of genes and the limited temporal resolution, we clustered genes based on the sign of expression changes between consecutive AS (+, −, or =), yielding 25 clusters with variable sizes, encompassing 1817 genes (**Fig. 5A**). Functional enrichment revealed a modular organization: clusters predominantly captured defense-associated secondary metabolism, carbon and structural reallocation, or transcriptional regulatory control, consistent with distinct but interconnected functional programs. Some clusters, such as + = − and − − =, highlighted tight coupling between metabolic and defense processes, whereas others, like + − +, suggested higher-order regulatory coordination (**SFig 1**). Collectively, these results demonstrated that repeated AS triggers structured and functionally coherent transcriptional programs that can be analyzed at the cluster level.

**Figure 5.**
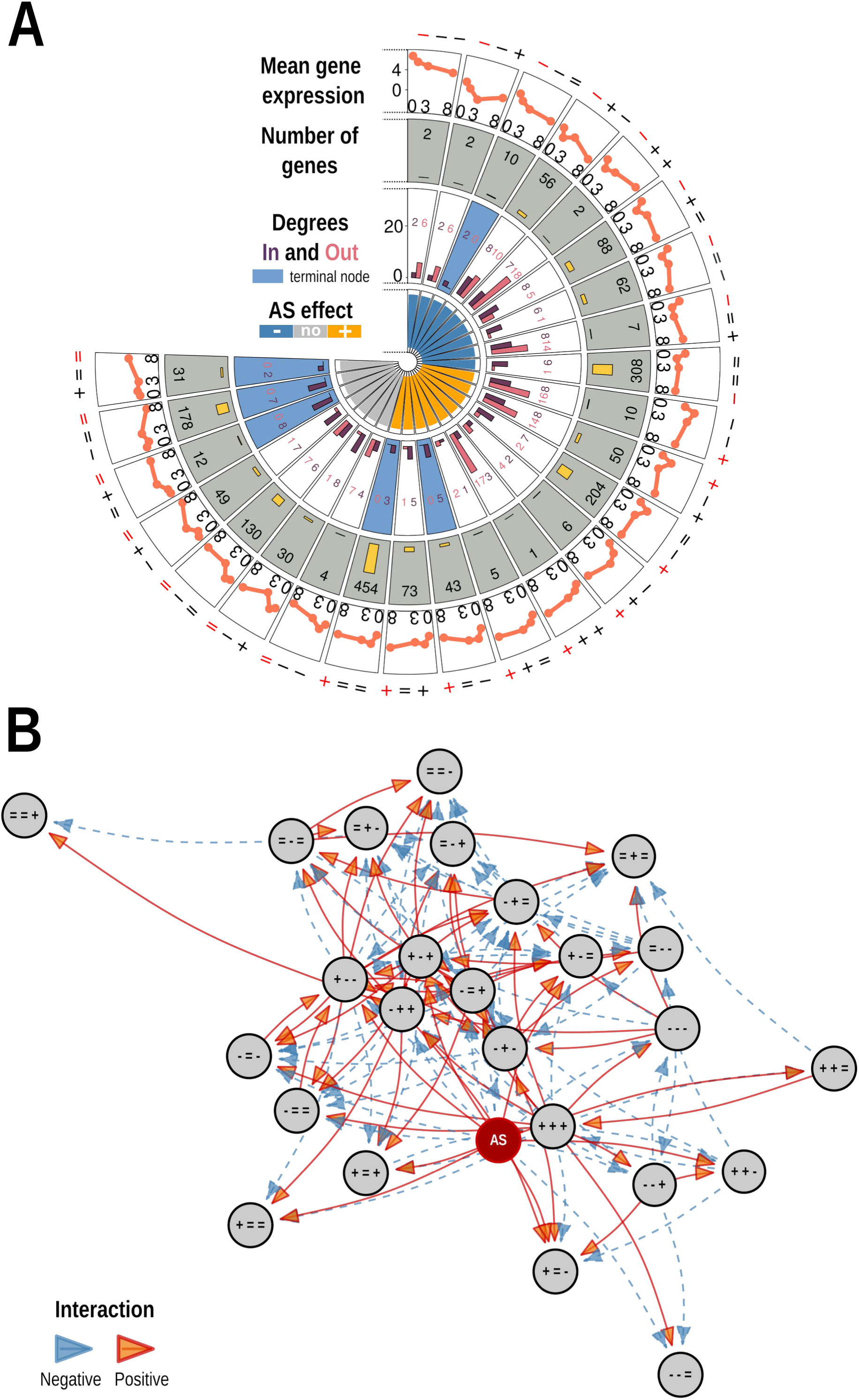
Dynamic model of plant priming through repeated acoustic stimulations (AS). **A.** Clustering of genes based on the sign of expression changes between consecutive acoustic stimulations (+, −, or =), defining 25 transcriptional clusters. The panel shows cluster size and their connectivity within the inferred network, as well as the direct effect of acoustic stimulation (AS_effect_) on each cluster. **B.** Inferred dynamic gene regulatory network at the cluster level. Nodes represent gene clusters and directed edges represent inferred regulatory influences between clusters. An additional node represents the direct effect of acoustic stimulation (AS_effect_).

To capture the modular organization underlying these coordinated transcriptional programs, we implemented a network modeling approach. Assuming time-independent molecular regulations, we described the dynamics of cluster-level gene expression using a linear iterative model of the form X_N+1_=A.X_N_+AS_effect_ where X_N_ denotes the representative expression levels for each gene cluster at the N^th^ AS. In this framework, A is a time-invariant regulation matrix that quantifies the influence of each cluster on all others, and AS_effect_ is a vector of fixed effects capturing the systematic impact of AS on cluster expression independently of inter-cluster regulation. Although this approach can point to candidate regulatory interactions between clusters, it should primarily be viewed as a descriptive framework that quantitatively characterizes the emergent dynamics of the system rather than as a direct inference of molecular causality. The inferred connectivity should therefore be interpreted as capturing dominant statistical dependencies rather than direct molecular regulations, making the model a quantitative description of regulatory-state transitions rather than a causal network.

The inferred dynamic network comprised 25 gene clusters with an additional node representing the AS effect and connected to 18 gene clusters (**Fig. 5B**). The network displayed an average node degree of 5.8 and a density of 0.23, corresponding to 151 directed edges. With a characteristic path length of 2.04, these metrics collectively indicated a “small-world” structure that was both compact and highly interconnected, despite a relatively low density. Six clusters behaved as terminal nodes, receiving regulatory inputs without outgoing connections, whereas the remaining 19 clusters exhibited both incoming and outgoing edges (**Fig. 5**). These clusters overall exhibited both incoming and outgoing edges. Together, this architecture revealed a densely connected, non-hierarchical regulatory organization rather than a strictly layered cascade, consistent with extensive cross-module interactions and distributed control of transcriptional dynamics.

**SFig 1.**
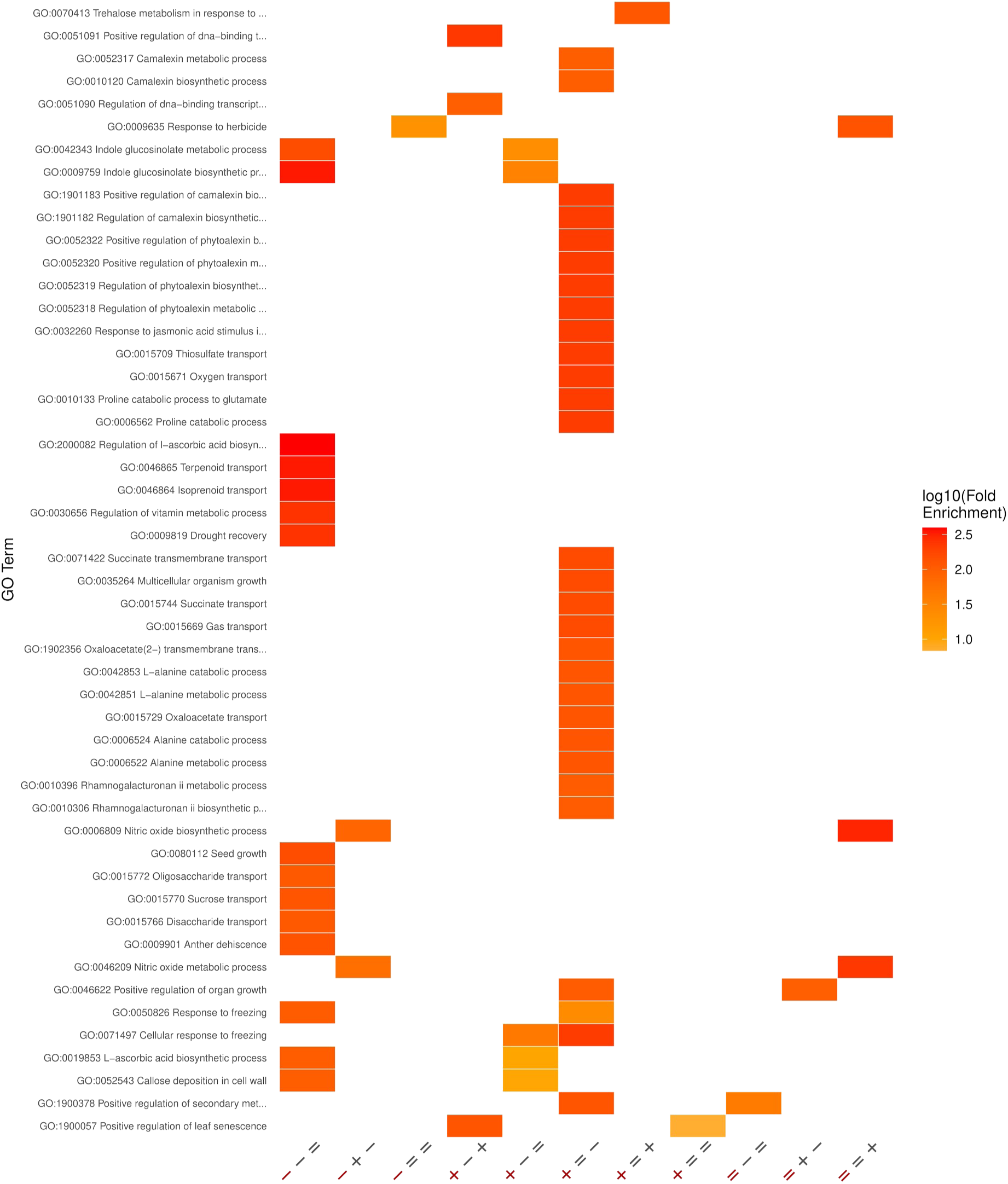
Functional enrichment of Gene Ontology biological process categories across transcriptionally defined gene clusters.

### Defense priming by AS is robust and accumulates progressively through near-neutral network modes

To determine how the topology of the transcriptional network structures gene regulatory dynamics, we performed a generalized modal decomposition of the inferred regulation matrix A. The matrix A exhibited 17 distinct eigenvalues, comprising 7 real eigenvalues and 10 forming complex-conjugate pairs (**Fig. 6A**). Notably, the eigenvalue λ = 1 had multiplicity 9, indicating the presence of a high-dimensional subspace of near-neutral modes in which perturbations were neither amplified nor strongly attenuated, thereby enabling persistent dynamical contributions to the system’s transcriptional response. This eigenvalue spectrum revealed three main dynamical regimes organized into 12 generalized modes (metamodes), which captured coordinated patterns of gene expression with distinct temporal behaviors. Among them, 6 metamodes (MM1–MM6) had eigenvalues with magnitude strictly below one, corresponding to stable components that decay following perturbation, either monotonically or through damped oscillations (**Fig. 6A**). One metamode (MM7) corresponded exactly to the eigenvalue λ = 1, defining neutral directions in which perturbations persisted over successive time steps without net amplification or decay (**Fig. 6A**). The remaining 5 metamodes (MM8–MM12) exhibited eigenvalues with magnitude greater than one, indicating dynamically amplifying directions in the transcriptional state space. The strongest amplification was observed for MM12 (λ = 1.48), while MM8 (λ ≈ 1.003) lay close to neutrality but retained a weakly amplifying behavior (**Fig. 6A**).

**Figure 6.**
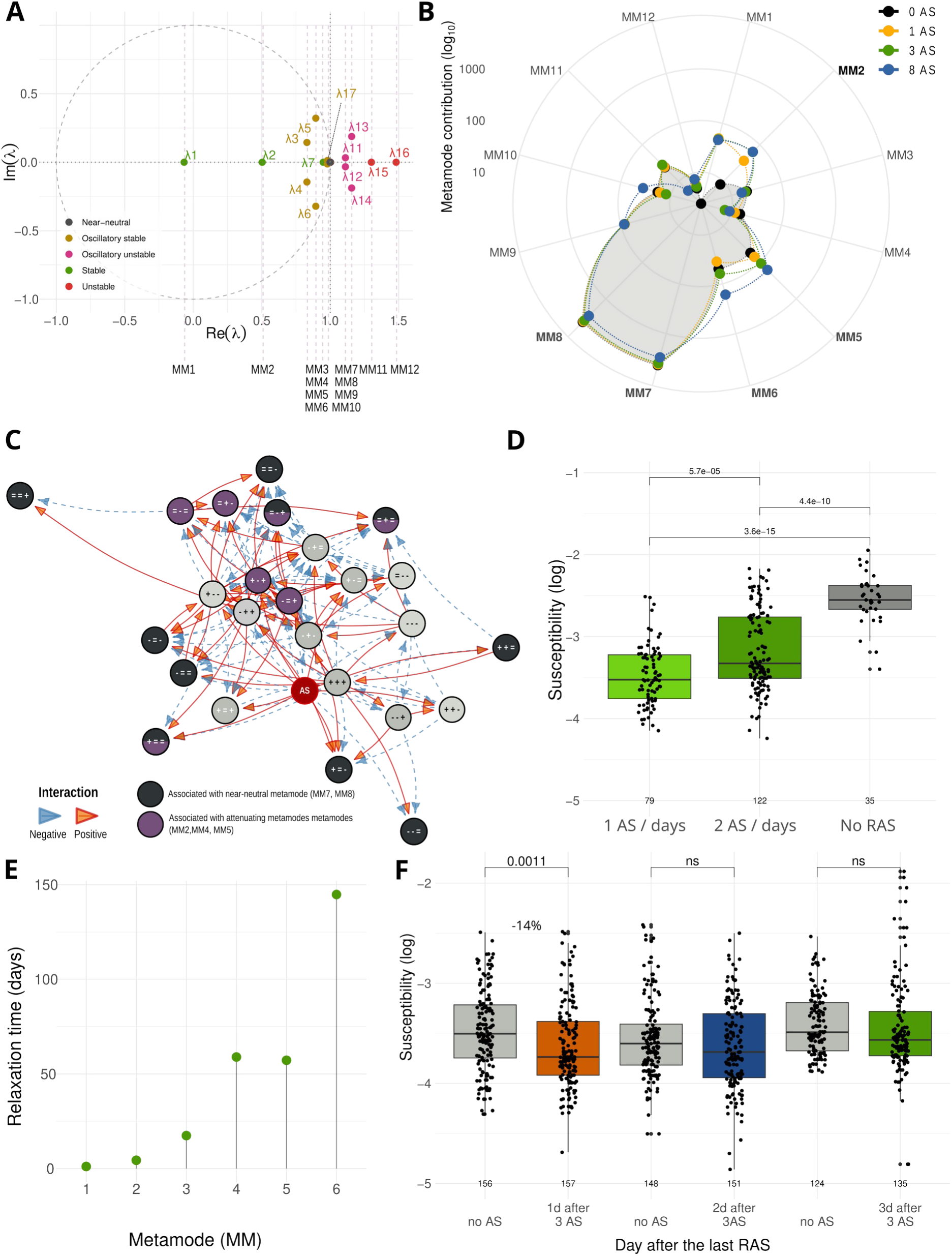
Network dynamics and robustness of plant defense priming induced by repeated acoustic stimulation (AS). **A.** Distribution of eigenvalues of the matrix A in the complex plane. Matrix A represents the inferred gene regulatory matrix describing how gene expression levels at time t influence those at the next time step. Eigenvalues located within the unit circle correspond to stable modes. The 17 eigenvalues formed 12 metamodes (MM) capturing dominant dynamical behaviors of the system. **B.** Projection of cluster-level gene expression onto the generalized modal decomposition, showing the contribution of each metamode to the transcriptional dynamics across repeated stimulations. **C.** Identification of network nodes (gene clusters) contributing to near-neutral and attenuating modes, highlighting distinct regulatory modules associated with slow (memory-retaining) versus fast (dissipative) dynamics. **D.** Phenotypic variation in susceptibility to *S. sclerotiorum* in plants exposed to increased daily frequency of AS. Boxplots represent the interquartile range (25th–75th percentile) with the median indicated by the bold line. Individual data points are overlaid using jitter for clarity. **E.** Characteristic relaxation times of stable metamodes, estimated from eigenvalues, indicating the temporal persistence of transcriptional perturbations following repeated AS (τ = −3 / log(|λ|). **F.** Phenotypic variations in susceptibility as a function of time after the initial three AS exposures.

To assess how these dynamical components contribute to stress memory induced by acoustic stimulation (AS), we projected gene-expression vectors corresponding to 0, 1, 3, and 8 AS exposures onto the modal basis and quantified modal amplitudes for each metamode (**Fig. 6B**). The near-neutral metamodes MM7 and MM8 (λ ≈ 1) dominated transcriptional dynamics under AS. Because perturbations along these directions were only weakly attenuated by the intrinsic network dynamics, repeated stimulations progressively accumulated within these modes, allowing them to integrate the history of prior acoustic exposures.

In contrast, attenuating metamodes such as MM4 (λ = 0.89), MM5 (λ = 0.94), and MM2 (λ = 0.50) showed increasing contributions with the number of stimulations, suggesting that dissipative components of the network also participated in shaping the integration and stabilization of transcriptional responses to repeated AS.

Each metamode was supported by regulatory subnetworks in which certain clusters played a more prominent role. Among the ten clusters contributing most strongly to the near-neutral metamodes MM7 and MM8, seven corresponded to terminal nodes of the regulatory network (**Fig. 6C**). These nodes integrated regulatory inputs from other clusters but did not regulate downstream clusters themselves (= = +, = = −, = − +, = + =, − = =, + = =, += −). Altogether, these clusters encompassed 1056 genes, indicating that the persistence of perturbations within near-neutral modes involves coordinated transcriptional responses across a large fraction of the transcriptome.

These clusters were not specific to a single dynamical regime. Three of the terminal clusters contributing to near-neutral modes were also involved in the attenuating metamodes MM4 and MM5. However, the additional clusters contributing to these attenuating modes corresponded primarily to intermediate nodes of the regulatory network involved in upstream regulatory interactions ( **Fig. 6C**). Their topological position was consistent with a stabilizing role, as perturbations reaching these nodes were redistributed through multiple regulatory interactions, including incoherent feed-forward loops, thereby promoting the dissipation and stabilization of transcriptional responses. As observed for the near-neutral modes, these stabilizing processes also involved several thousand genes. Together, these observations indicated that stress memory did not appear to arise from the persistent regulation of a small number of individual genes, but rather emerged from the collective dynamics of large regulatory subnetworks shaping the global transcriptional state.

These dynamical interactions between regulatory clusters defined the intrinsic response times of the system and generated experimentally testable predictions regarding both the formation and decay of stress memory. As described above, the establishment of memory was associated with the near-neutral metamodes MM7 and MM8, which accumulated perturbations only slowly and therefore constrained the rate at which stress memory could form. Consequently, increasing the frequency of acoustic stimulation (AS) exposures was not expected to substantially accelerate the emergence of the resistance phenotype. Experimental tests confirmed this prediction. Increasing the stimulation frequency to twice daily over three days did not further enhance resistance; instead, it led to a slight reduction compared to the once-daily regime (**Fig. 6D**). Such a response confirms that the transcriptional network integrates repeated cues progressively rather than instantaneously.

The decay of stress memory could similarly be interpreted in terms of the stable components of the network dynamics. The attenuating metamodes defined characteristic relaxation times according to τ = −1/log(|λ|). For the metamodes most strongly activated by AS, three characteristic times (3τ) ranged from 1.10 days for MM1 to 44.80 days for MM6 (**Fig. 6E**).

To experimentally test this prediction, Col-0 plants were subjected to three repeated AS treatments and inoculated with *Sclerotinia sclerotiorum* 0, 1, 2, or 3 days after the last stimulation. Naïve plants (no AS) were inoculated in parallel as controls (**Fig. 6F**). Inoculation immediately after three AS treatments led to a 25% reduction in susceptibility compared with naïve plants (**Fig. 3B**). When inoculation was delayed by one day after the last stimulation, susceptibility remained reduced by 14% (p = 6.2 × 10⁻³; **Fig. 6F**). However, the priming effect disappeared when inoculation occurred two or more days after the last stimulation (p = 0.23 at 2 days and p = 0.99 at 3 days; **Fig. 6F**). Consistent with the predictions of the dynamical model, the defense priming effect induced by AS therefore vanished between one and two days after the cessation of stimulation.

Taken together, these results demonstrated that acoustic stimulation shaped plant defense through a dynamical process in which network-encoded stress memory emerged from the interplay between slowly accumulating near-neutral modes and dissipative stabilizing modes of the regulatory network. In this framework, repeated acoustic stimuli progressively drove the transcriptional system from a naïve toward a primed state, while the intrinsic relaxation dynamics of stable modes determined the limited temporal persistence of this memory. These results therefore provided a mechanistic framework linking the topology of transcriptional regulatory networks to the dynamics of sound-induced defense priming.

## Discussion

Stress memory is closely associated with the priming of plant defense responses, yet the mechanisms underlying the transition from a naïve to a primed state remain poorly understood. In this context, priming is defined as an acquired state in which prior stimulation enhances the magnitude and/or speed of the response to a subsequent stress. Here, we show that this state emerges from the dynamic reconfiguration of the transcriptional network, which allows the coordinated mobilization of multiple molecular processes, rather than from the sustained activation of specific genes. In this study, we demonstrate that repeated acoustic stimulation (AS) induces a transient but robust priming of quantitative disease resistance (QDR) in *Arabidopsis thaliana*. Three consecutive stimulations were sufficient to maximize resistance, which then plateaued despite additional exposures and progressively decayed within two days after stimulation ceased.

Our results indicate that this priming effect is tightly linked to the establishment of a network-encoded stress memory. In non-inoculated plants, repeated AS induced the expression of defense-related genes, while upon infection, it enabled the recruitment of a large set of additional genes and amplified the response of thousands of inoculation-responsive genes. Importantly, this response was not driven by a small set of key regulators but instead emerged from coordinated changes across a large number of genes. Consistent with this, our network analysis revealed a densely connected and non-hierarchical regulatory structure, suggesting that priming arises from distributed interactions rather than linear signaling cascades. Nevertheless, establishing a direct causal link between transcriptome-wide reprogramming and the observed phenotypic resistance gains remains challenging. Non-transcriptional mechanisms — including post-translational modifications, metabolite accumulation, and cell wall compositional changes — may contribute independently to the resistance phenotype and warrant investigation in future work.

Interestingly, while a single acoustic stimulation (1 AS) activated a core set of defense-related genes, it transiently increased plant susceptibility to *S. sclerotiorum*. This apparent contradiction may reflect an initial metabolic cost or a temporary trade-off before the transcriptional network converges toward a coordinated and effective immune response. Our dynamic model supports this view, as stabilizing and attenuating metamodes (e.g., MM1, MM2) require repeated exposures to fully deploy their homeostatic and protective regulatory functions.

The dynamic modeling further provides a mechanistic framework for this behavior. The coexistence of slowly relaxing (near-neutral) and rapidly dissipating modes offers a simple explanation for both the accumulation and the decay of stress memory. Repeated stimulations progressively reinforce slow modes, leading to enhanced responsiveness, while the absence of stimulation allows the system to relax back toward its basal state. This dynamic explains both the saturation of resistance after three stimulations and its transient nature over time. In this context, priming can be viewed as an emergent property of the transcriptional network, arising from its intrinsic dynamical structure which coordinates and integrates individual regulatory events, rather than operating in isolation from specific gene activations.

Epigenetic regulation may also contribute to the establishment and maintenance of stress memory, potentially by influencing the regulatory states and dynamics captured at the transcriptional level. A subset of known stress memory markers was modulated by AS, consistent with enrichment analyses and supporting a potential contribution of chromatin-based mechanisms to priming. In parallel, we observed signatures consistent with increased RNA turnover, which may facilitate rapid transcriptome reconfiguration. Together, these processes could enable both the stabilization of memory during repeated stimulation and its reversibility once stimulation ceases. The relatively short duration of the primed state further suggests that AS-induced memory does not rely on long-term structural modifications, but rather on dynamic and reversible regulatory processes. Our results reveal that network-level regulation can encode a distributed stress memory, while robust to genetic perturbations, is subject to intrinsic dynamical constraints that limit both the magnitude and duration of acoustic priming. Whether transcriptional priming can overcome these intrinsic limits without additional stabilizing mechanisms remains an important question for crop protection.

Although the dynamic model captures key features of the transcriptional dynamics associated with stress memory and predicts phenotypic outcomes, it does not imply that transcriptional regulation constitutes the sole molecular substrate of memory. The inferred interactions reflect statistical dependencies between gene clusters rather than direct molecular regulations. Increasing the temporal resolution of transcriptomic measurements would likely improve the identification of early signaling events and provide deeper insight into the mechanisms by which acoustic stimuli are perceived and transduced.

More broadly, our findings support the emerging view that acoustic signals may represent a relevant component of the plant sensory environment. These findings should be interpreted in the context of a growing literature suggesting beneficial effects of acoustic stimulation on plant health. However, many of these studies rely on the expression of a limited number of marker genes as proxies for defense activation, without directly quantifying disease resistance. By pairing genome-scale transcriptomics with phenotypic assays, we demonstrate that the link between acoustic-induced gene expression and immunity is highly nuanced, with the magnitude and durability of resistance gains being bound to intrinsic dynamical constraints. We therefore propose that future studies in this field would benefit from integrating direct resistance assays alongside molecular analyses, to provide a more complete picture of the agronomic relevance of acoustic stimulation. Although our experiments were restricted to a single frequency, they demonstrate that repeated acoustic stimulation induces a measurable but transient priming effect, whose biological significance needs to be evaluated in the context of the dynamical constraints identified here.

## Conclusion

In this study, we investigated how repeated acoustic stimulations prime the plant immune response. A key finding is that this stress memory does not simply rely on standalone molecular switches; rather, it emerges from a distributed regulatory architecture where individual regulatory events are integrated into the collective dynamics of thousands of genes. This distributed architecture explains both the robustness of priming — its independence from individual gene perturbations — and its intrinsic limits, namely the rapid decay of memory and the saturation of resistance gains. RNA-seq analyses revealed that these stimulations dynamically reshape the transcriptome, modulating a substantial fraction of genes in *Arabidopsis thaliana* over time. To capture this complexity, we inferred a dynamic genome-scale gene network, which revealed how network-level regulatory structure governs the robustness and temporal constraints of the system. The short-lived nature of this memory suggests that longer-lasting priming may require additional mechanisms, including epigenetic regulation. Importantly, our work demonstrates that adopting a systems-level perspective on transcriptional dynamics provides a richer understanding of priming, showing how the coordinated activity of thousands of genes drives robust yet transient defense responses. Beyond its implications for understanding stress memory as an emergent property of regulatory-network dynamics, our findings provide a framework for understanding how repeated mechanical or environmental stimuli can progressively reshape plant defense states over time.

## Data availability

To review GEO accession GSE293061:

Go to https://www.ncbi.nlm.nih.gov/geo/query/acc.cgi?acc=GSE293061 Enter token ojcnwuaghxuxxup into the box

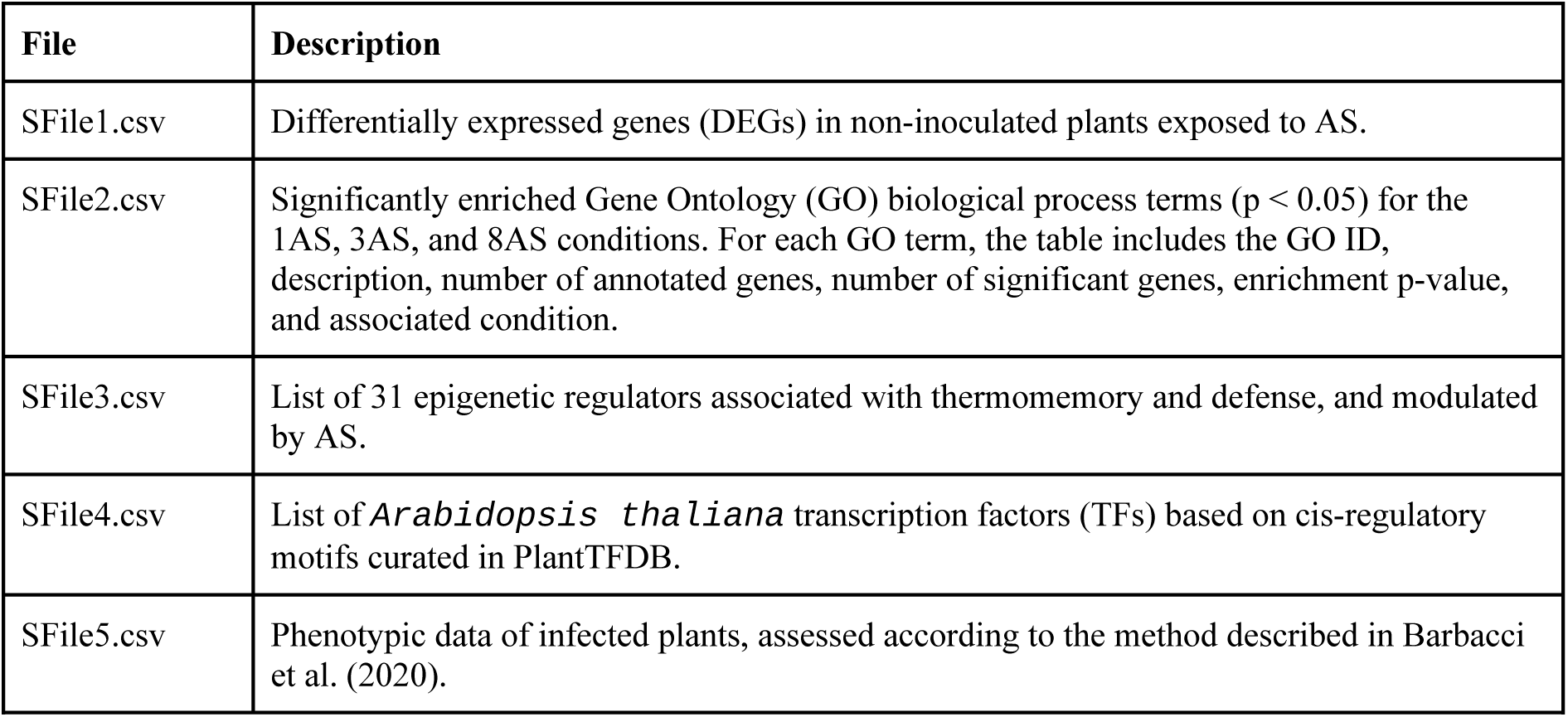
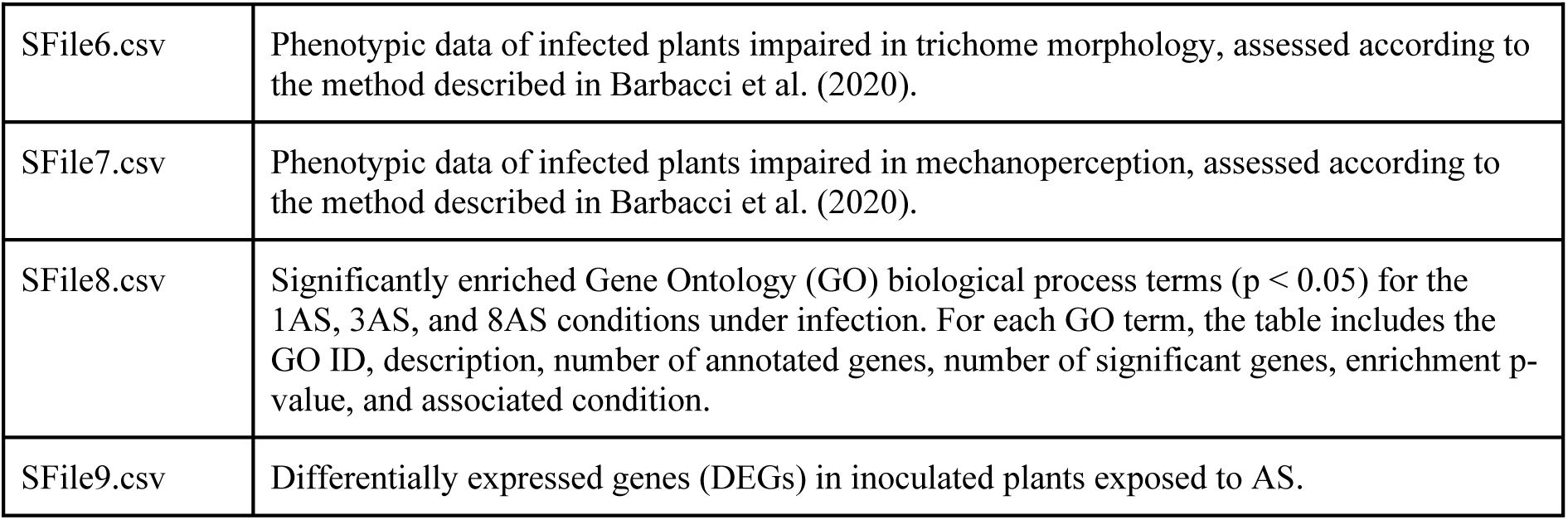

## Acknowledgment

Khaoula Hadj-Amor’s thesis was funded by INRAe’s MathNumm department and the Occitanie region. AB and FG are funded by the templates project of the DIGITBIO metaprogram, the Thigmoimmunity project of the INRAe plant health department, and the Explorae France 2030 program SAPIENS. SR was funded by the ERC VaryWhim. The authors would like to thank Jean-Marie Frachisse and Nathalie Leblanc-Fournier for providing seeds

## Author contributions

AB, AD, FG, KHA and SR planned and designed the research. AD designed the experimental device. OL, MK and PCS performed the experiments, including phenotyping and RNA-seq sampling. KHA designed the mathematical model and the clustering. AB, FG, KHA and SR analyzed and interpreted the RNA-seq data and modeling results. MK conducted the phenotypic validation of the model. AB and KHA prepared the figures. AB wrote the manuscript. AB, FG, KHA and SR reviewed and improved the manuscript, with input and approval from all authors. OL and SR contributed equally to this work. AB and FG contributed equally to this work.

## Declaration of generative AI and AI-assisted technologies in the manuscript preparation process

During the preparation of this work the author(s) used ChatGPT to assist with language editing of the manuscript text. After using this tool/service, the author(s) reviewed and edited the content as needed and take full responsibility for the content of the published article.

